# Lipid composition and mechanical force underlie multi-modal regulation of Piezo1 gating

**DOI:** 10.1101/2025.10.24.684433

**Authors:** George Vaisey, Roderick MacKinnon

## Abstract

Piezo1 ion channels are widely expressed cellular mechanosensors. They adopt an intrinsically curved shape when closed and are thought to open when mechanical forces applied to the membrane favor a more flattened conformation. By reconstituting Piezo1 channels into lipid vesicles, a flattened conformation has been determined, however, the ion conduction pore remained closed. In line with this observation, Piezo1 channels do not exhibit mechanical activation in the kind of lipids used in the structural studies. Here we show first that Piezo1 channels in cell-derived membranes retain functional mechanical gating, and second, that in cell-derived membranes they adopt a completely flattened disk shape associated with large conformational changes within and around the ion conduction pathway. These conformational changes occurring in cell-derived lipid membranes, suggest that mechanical force is necessary but insufficient, and that a specific membrane-derived cofactor apparently complements mechanical force to activate Piezo1.

---

Many living cells depend on rapid responses to mechanical cues. Piezo ion channels are on the frontline of this sensory process in mammalian cells (*1–4*). Structural studies of Piezo1 in detergent micelles or in liposomes show that it forms a trimeric assembly that is intrinsically curved under resting conditions, but that it can be flattened under applied force(*5–9*). If flattening were coupled to pore opening, as the current model of Piezo1 mechanical gating posits, we would have a simple description of mechanical gating whereby lateral membrane tension would open the pore by favoring the flattened, in-plane expanded conformation. Unfortunately, an important piece of this explanation is missing, because so far a flattened structure does not exhibit an open pore. Perhaps related to this missing piece is the finding that Piezo1 fails to activate, i.e., open functionally, in membranes comprising the kind of lipids used in structural studies.

Compared to other mechanosensitive channels, including the prokaryotic MSCs or eukaryotic two-pore-potassium channels (K2Ps), Piezo channels have in general remained refractory to robust functional studies in reconstituted systems(*10–14*). Spontaneous Piezo1 channel activity has been reported in lipid droplet bilayers and membrane patches of proteoliposomes(*15*, *16*) but reversible mechanically-activated currents have not been demonstrated and the measured single channel conductance is inconsistent with patch-clamp recordings of endogenous and overexpressed Piezo1(*1*, *17–21*). Here, we show first that Piezo1 ion channel activity is maintained in vesicles released from N-Ethylmaleimide (NEM) treatment of cells. Then we determine structures of Piezo1 channels in vesicles derived from these same membranes under varying membrane bending forces applied through the curved vesicle membrane. The structures reveal conformational changes that appear to map directly to channel activation and imply a multi-modal basis for Piezo1 channel gating.

## Results

### Cell membrane-derived vesicles retain Piezo1 channel function

We first reconstituted Piezo1 channels into small unilamellar vesicles (SUVs) consisting of defined lipid compositions used in past structural analyses of Piezo1 in membranes(*8*) (fig. S1). From the SUVs we generated giant unilamellar vesicles (GUVs), microns in size, using established protocols(*22*, *23*) and confirmed incorporation of Piezo1 by detecting the fluorescence of a C-terminally fused GFP (Fig. 1A). Even though Piezo1 channels were present in the membrane, we could not detect channel activity by patch recording (Fig. 1B). We varied Piezo1 purification conditions, lipid composition, and conditions for forming GUVs, all to no avail regarding channel activity. We occasionally observed transient ionic currents of varying amplitudes that were not reproducibly mechanically activated by patch pressurization (Fig. 1B). As we have found in other studies, reconstitution of membrane proteins at high concentrations can yield this kind of channel-like activity(*24*), but without the correct single channel conductance and mechanical activation, these do not represent bona fide ion channel recordings.

**Figure 1.**
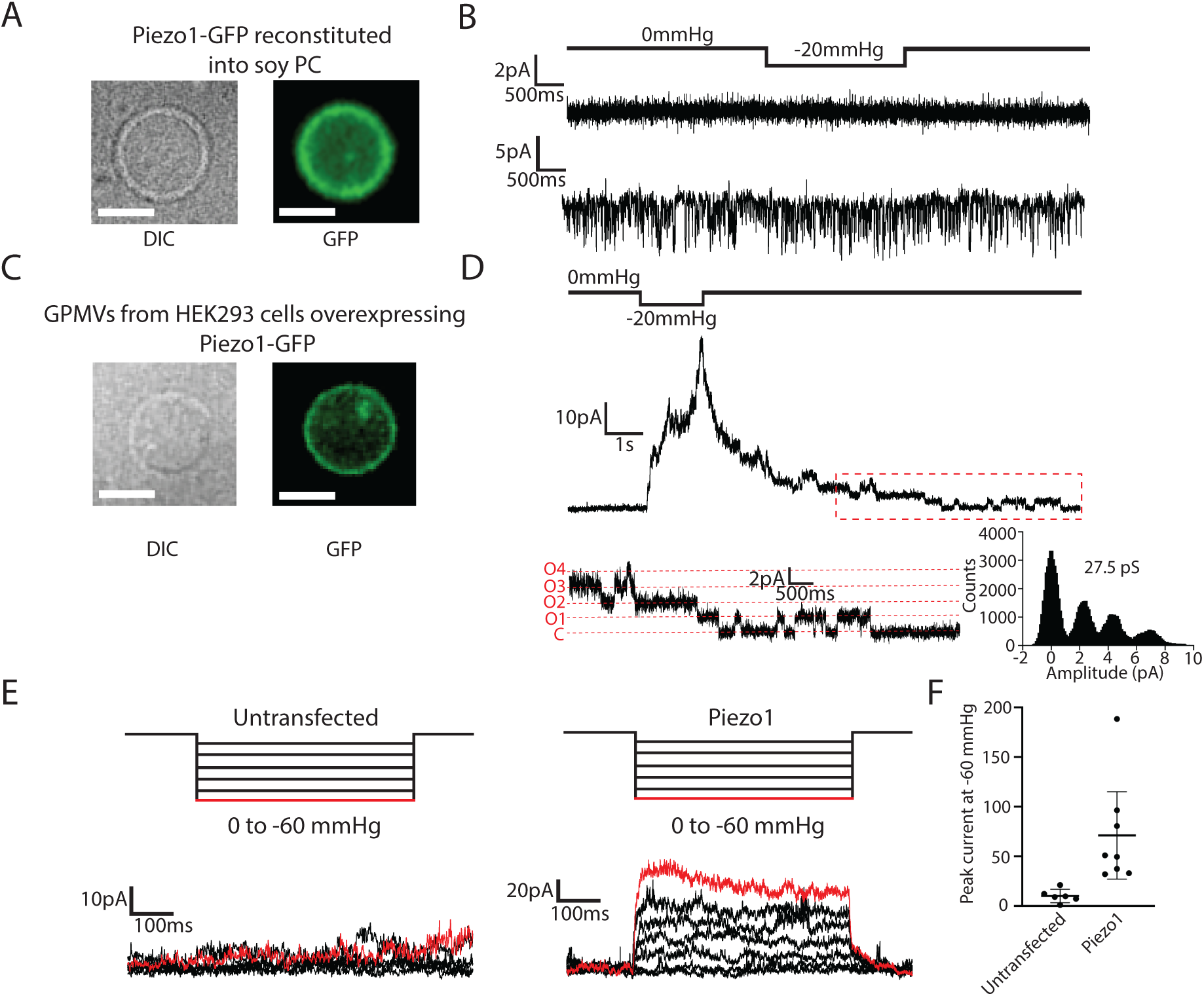
Functional mechanical activation of Piezo1 in GPMVs but not when reconstituted into soy PC liposomes. (A) DIC (left) and fluorescence image (right) of purified Piezo1-GFP reconstituted into soy PC liposomes that were then swelled into giant unilamellar vesicles. Scale bar 10μm. (B) Two example patch-clamp recordings of patches excised from GUVs as prepared in *A*. Recordings are made holding at -80mV with a 2s pulse of -20mmHg. (C) Images taken as in *A* of giant plasma membrane vesicles (GPMVs) generated by N-ethylmaleimide (NEM) incubation of cells overexpressing Piezo1-GFP. (D) Electrical recording of a patch excised from a GPMV overexpressing Piezo1-GFP. The patch is held at +80 mV and a 1.5 s pressure pulse at - 20 mmHg is made to elicit Piezo1 channel openings. The red-dashed inset shows a 5s duration of single-channel openings and on the right the corresponding amplitude histogram is shown, giving a single channel conductance of 27.5 pS. (E) Excised patch recordings of GPMVs from Piezo1 knockout HEK293 cells that were either untransfected or overexpressing Piezo1. Patches were excised and held at +60mV with increasing -10mmHg pressure steps up to -60 mmHg. (F) Data from recordings as in *E* plotted for untransfected (n=6) or Piezo1-overxpressing (n=8) GPMVs.

Exploring alternative approaches, we experimented with the formation of plasma membrane vesicles (PMVs) by incubation of cells with N-ethylmaleimide (NEM) in the presence of millimolar free calcium(*25*, *26*). Adherent HEK293 cells treated with NEM generate giant PMVs (GPMVs), as shown in Figure 1C for cells overexpressing mPiezo1-GFP. Electrical recordings of membrane patches excised from GPMVs yielded reversibly pressure-activated currents. When channels were few enough to detect single channel events, we measured the single channel conductance to be about 27.5 pS (Fig. 1D), consistent with cellular recordings of Piezo1 by us and others(*1*, *17–21*). Macroscopic recordings of GPMVs derived from cells overexpressing Piezo1 showed increases in current in response to stepwise pressure pulses, with a mean peak current amplitude at -60 mmHg of 71 ± 49 pA recorded at +60 mV (Fig. 1 E and F). By contrast, we did not observe significant pressure-activated currents in GPMVs from Piezo1 knockout cells (Fig. 1 E and F). We note that the Piezo1 currents in GPMVs do not inactivate like in cells, however, with respect to single channel conductance and mechanical activation, the currents are very similar to Piezo1 channels in cells. We next examined whether Piezo1 structures in cell-derived membranes, which support function, appear different than Piezo1 structures in reconstituted membranes, which do not support function.

### Determination of outside-in and outside-out Piezo1 structures in cell-derived membranes

To determine Piezo1 structures in vesicles derived from the PMVs we used an approach developed in our laboratory to study other ion channel structures(*27*), but with modifications to overcome lower levels of Piezo1 expression and to selectively bias preparations towards outside-out and outside-in orientations of the Piezo1 channel (Methods). In brief, after NEM treatment of suspension HEK293 cells, centrifugation was used to separate the GPMVs from what we call small plasma membrane vesicles (SPMVs), whose size is reduced further by gentle sonication for cryo-electron microscopy (cryo-EM). After observing that the SPMVs contained Piezo1 in both outside-in and outside-out orientations in electron micrographs, we used affinity purification methods targeting either an intracellular GFP tag or an extracellular ALFA tag to separately enrich the two populations of channels for single particle cryo-EM data collection and processing. In both cases, the final preparation of SPMVs showed a dominant Piezo1 band on SDS-PAGE after Coomassie staining (Fig. 2A). SPMVs with Piezo1 adopting the outside-in orientation exhibited a tear-drop shape owing to membrane distortion created by the curved Piezo1 channel, as was reported previously(*6*, *8*, *28*, *29*) (Fig. 2A). Processing of the outside-in dataset using SPMVs in the size range 11 to 20 nm vesicle radius, yielded a structure at 3.7Å resolution (fig. S2 and S3). This resembled the curved structures of Piezo1 previously determined in detergent micelles and in liposomes(*5*, *6*, *8*), with a mean radius of curvature of about 11.8 nm (Fig. 2B and fig. S5). The refined structural model contains 1352 residues (out of 2547), with the first four transmembrane helical unit (THU) repeats and flexible loops missing. We note that this structure, like outside-in structures in small vesicles determined before, is highly curved.

**Figure 2.**
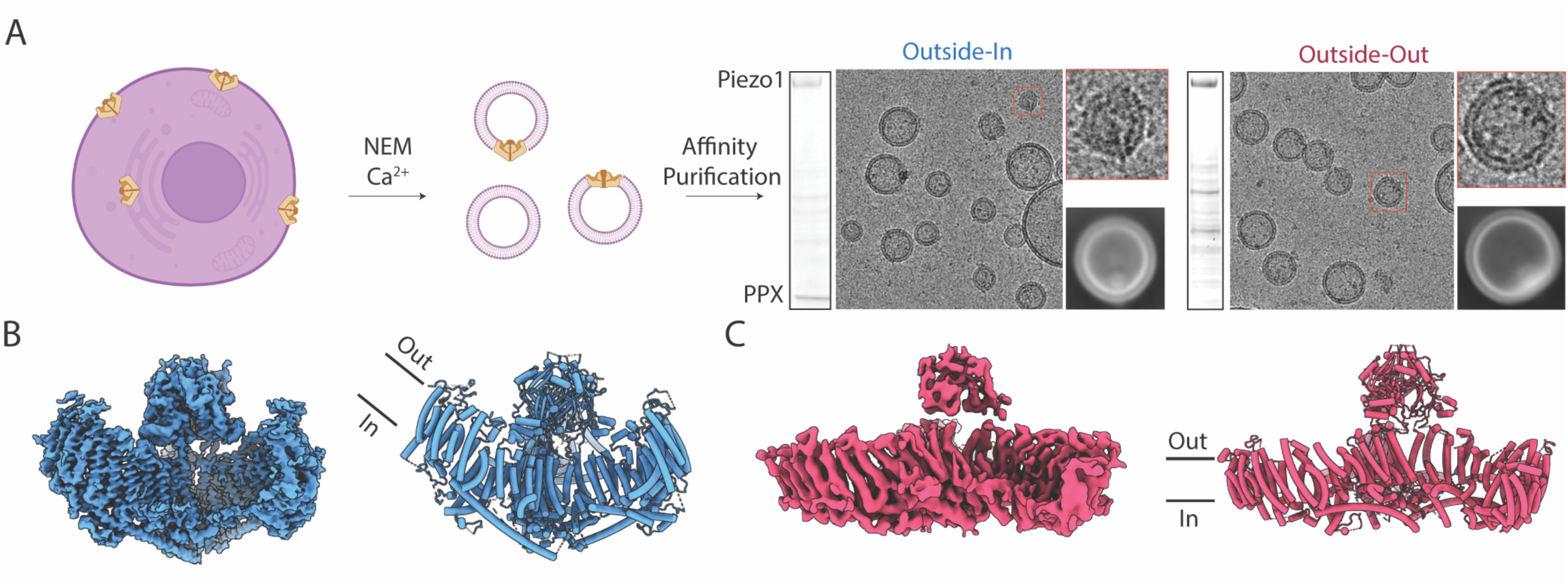
Methodology to isolate outside-in and outside-out Piezo1 in PMVs and determine single particle cryo-EM structures. (A) (left) Schematic illustrating basic approach of obtaining Piezo1-containing PMVs. (right) Coomassie stained denaturing gels and representative micrograph images of the final outside-in and outside-out Piezo1 PMV samples, along with a representative 2D class image. (B) Cryo-EM map (left) and structure of outside-in Piezo1 in PMVs. (C) Cryo-EM map (left) and structure of outside-out Piezo1 in PMVs.

Outside-out Piezo1 SPMVs by contrast exhibit a different shape on micrographs, where the opposing curvature of the vesicle relative to the intrinsic curvature of Piezo1 produces a bending force that completely flattens the channel into a conformation in which the transmembrane arms are coplanar to the bilayer (Fig. 2A and C, fig. S4 and fig. S5). Despite similar sample quality and data acquisition settings to our outside-in map, the outside-out map reaches only a resolution of ∼6 Å (fig. S3). Even at this lower resolution, the map defined a region of the protein with sufficient detail to build a C*α* model extending from residue 589 to 2547 (Fig. 2C).

### Conformational differences between curved and flat Piezo1 channels in cell-derived membranes

Figure 3A shows the outside-in and outside-out models of Piezo1 superimposed. As shown previously, Piezo1’s shape is highly dependent on membrane bending forces(*28*, *29*). The outside-in structure is curved. This is because the channel tends to follow the intrinsic curvature of the vesicle. But we know that Piezo1 does not conform perfectly to the vesicle shape, which is why outside-in Piezo1 vesicles are tear drop shaped rather than spherical(*28*, *29*) (Fig. 4A). The shape of the Piezo1 vesicle reflects the energy minimum condition reached when forces between the vesicle membrane and Piezo1 (including the lipid membrane between its three protein arms, called the Piezo dome) exactly counterbalance each other. Consequently, Piezo1 is more curved than it would be in a planar membrane, and the vesicle deviates from a sphere. Similarly, the shape adopted by Piezo1 in an outside-out orientation must also reflect an energy minimum configuration. In this case, we observe a flat, disk-shaped Piezo1 channel and a vesicle that is distorted towards an oblate spheroid (Fig. 4D).

**Figure 3.**
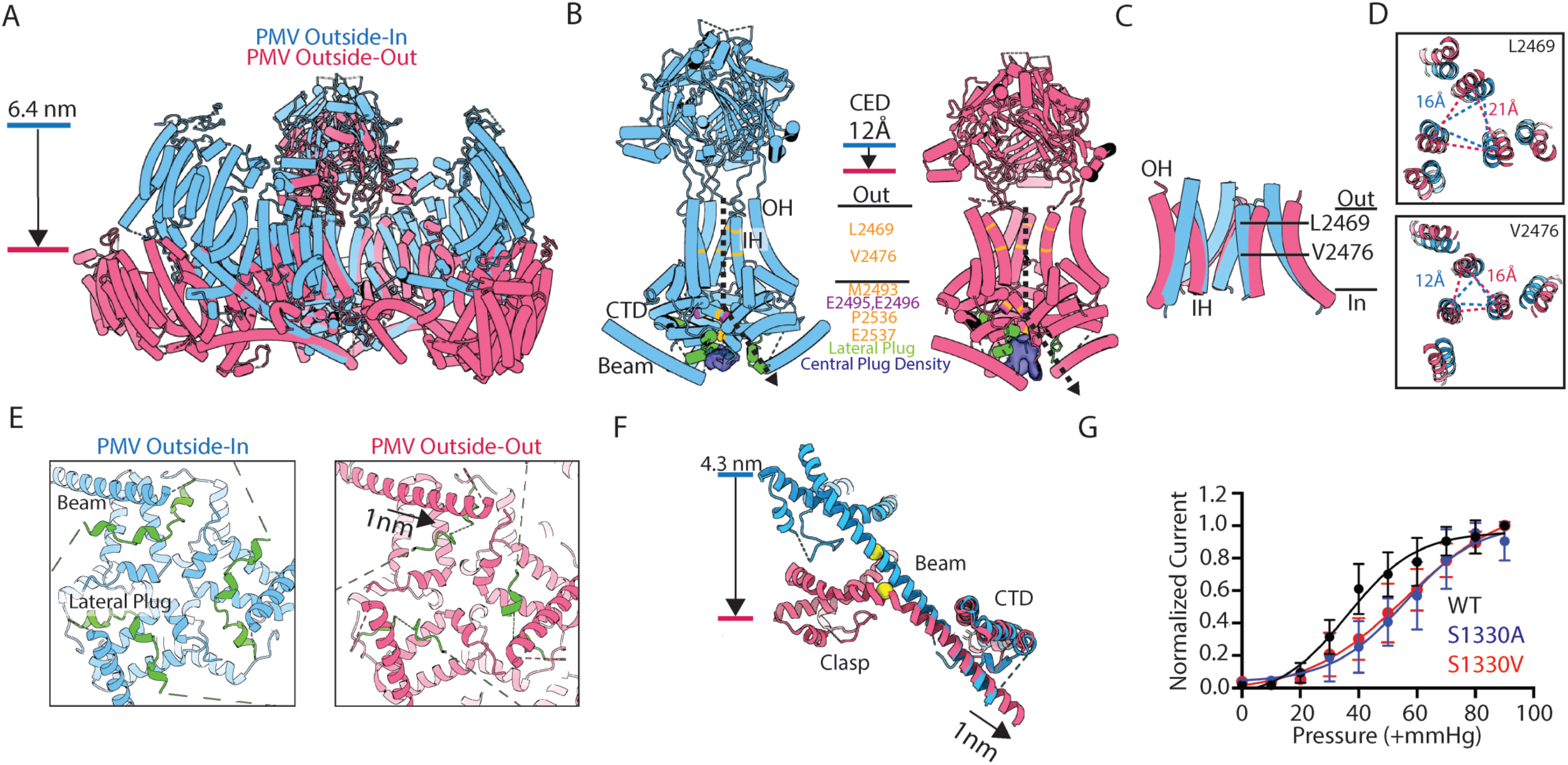
Conformational differences between curved and flat Piezo1 channels in cell-derived membranes. (A) Aligned structures of outside-in (blue) and outside-out (red) oriented Piezo1 from PMVs. The major displacement of the transmembrane arms is highlighted. (B) Isolated views of regions of Piezo1 involved in channel gating and ion conduction for the two Piezo1 structures are shown. The downward displacement of the CED as Piezo1 flattens is annotated. Residues that form hydrophobic constrictions are color-coded as are residues identified as important for ion conduction. The lateral plug is highlighted and cryo-EM density for the central plug density is shown. The dashed arrow indicates the proposed ion conduction pathway for Piezo1. (C) Isolated view of the aligned OH and IH for the two Piezo1 structures. (D) Top-down views of two residue positions (L2469 and V2476) that form part of the hydrophobic constriction in the transmembrane pore of Piezo1. The distance between the C*α* atoms of neighboring subunits is measured and annotated. (E) Isolated views of the cytosolic side of Piezo1 at the three-fold axis. The beam helix and lateral plug (green) are annotated. The 1 nm sliding of the beam helix towards the central three-fold axis appears to displace the lateral plug in the outside-out oriented Piezo1 structure. (F) An isolated view of the Piezo1 beam helix, clasp and CTD region from the aligned outside-out and outside-in PMV structures. Serine residue 1330, positioned proximal to a helical kink in the outside-out PMV Piezo1 structure, is depicted as yellow spheres in the two structures. (G) Outside-out pressure clamp recordings were performed on cells overexpressing WT, S1330A or S1330V Piezo1 variants. Currents were held at +60mV and pressurized stepwise from 0 to -90mmHg. Peak currents at each pressure step were normalized to the peak current at -90mmHg and the datapoints were fit to a sigmoidal curve. P_50_ values are 36 ± 5mmHg (WT), 56 ± 6mmHg (S1330A) and 55 ± 6mmHg (S1330V). N=5 for all conditions.

**Figure 4.**
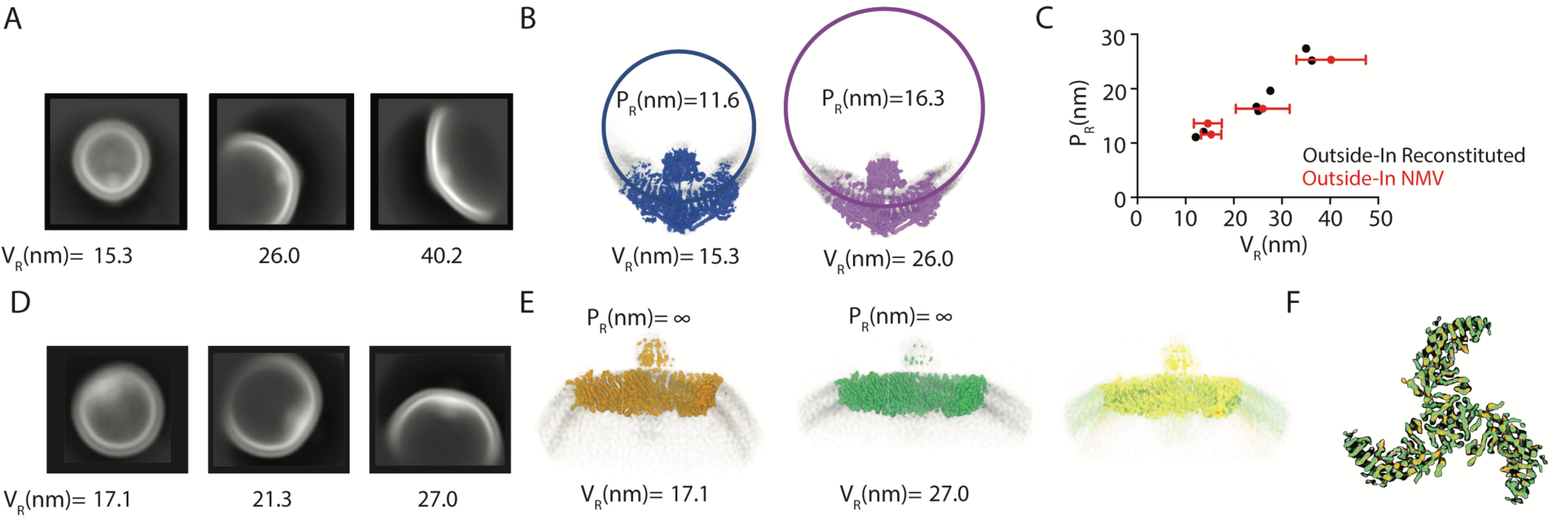
Piezo1 shape change analysis in PMVs of different size. (A) Example images of 3 populations of 2D classes generated by classification of aligned Piezo1 particles after extraction with a 60 nm box size. The mean vesicle radius, V_R_ is given below each image. (B) Two example single particle cryo-EM maps of Piezo1 from different vesicle size classes are shown in blue and purple. A circle is fit to the mid-plane of the Piezo1 transmembrane helices to estimate the Piezo1 radius of curvature (P_R_) which is annotated along with the mean vesicle radius the particles came from. (C) P_R_ is plotted as a function of V_R_ for 4 classes of outside-in PMV Piezo1 structures, where V_R_ is the mean value derived from measurements of 30 vesicles for each class of particles and the error bar represents the standard deviation. Overlaid are previously published values from tomographic analysis of Piezo1 reconstituted into liposomes in the outside-in orientation^8^. (D) As in *A* but here example images are given for Piezo1 particles from the outside-out PMV dataset after extraction with an 80 nm box size. (E) Two example single particle cryo-EM maps of Piezo1 from the smallest and largest vesicle size classes in the outside-out PMV dataset are given in orange and green. On the right an overlay of these two maps shows that the two structures are essentially identical. (F) A top-down view of the two outside-out Piezo1 structures as in *E*, showing identical conformations.

What local conformational changes occur inside the Piezo1 protein when it goes from highly curved to flat? In figure 3B we show isolated structural elements thought to be important for channel gating and forming the ion conduction pore of Piezo1. These elements include the cytoplasmic extracellular domain (CED), outer and inner-helices (OH and IH) that form the transmembrane pore, and the C-terminal domain (CTD) that forms the intracellular region of the Piezo1 pore(*19*, *30*, *31*). In the flattened conformation relative to the curved, the CED is displaced ∼12Å towards the IH and OH (Fig. 3B and movie 1). This conformational change is notable because cysteine cross-links expected to limit the mobility of the CED inhibit mechanical gating(*32*). In addition, the OH and IH, which are attached directly to the CED via short linkers, are expanded in the flattened structure so that the pore’s extracellular vestibule is dilated (Fig. 3B and C). Further into the membrane along the channel’s 3-fold axis the distance between C*α* atoms of amino acids L2469 and V2476 are increased by ∼ 5 Å and ∼4 Å, respectively in the flattened conformation (Fig. 3C). These specific amino acids have been identified through mutagenesis studies as forming a hydrophobic gate^31,33^.

Numerous mutations have identified amino acids that affect ion conduction(*19*, *31*, *33*, *34*) and define what is thought to be the ion conduction pore (Fig. 3B). The resolution of our structures is too low to define the pore’s chemistry, but conformational differences along the pore are evident when comparing the curved and flat structures. In addition to the changes already mentioned involving the CED and dilation along the 3-fold axis, we also observe differences near the cytoplasmic opening (Fig. 3E). At the cytosolic end of the 3-fold axis narrow constrictions are formed by residues M2493, P2536 and E2537, which would prevent ion conduction (Fig. 3B). Instead, structural and mutagenesis studies support the idea that the pore trifurcates laterally into portals as it approaches the cytoplasm. In particular, mutation of two highly conserved glutamate residues, E2495 and E2496, which line the lateral portal, alters ion permeability and conductance(*33*). In all previous structural studies, whether curved or flattened, the lateral portals are occluded by the lateral plug(*5–8*). The lateral plug is well-resolved in the curved structure but in the flattened structure the associated cryo-EM density appears fragmented and displaced, leading to the appearance of a cavity that might permit ion conduction (Fig. 3E fig. S6).

In addition to the structural changes that appear to open the pore in the flat structure, we highlight in Figure 3F structural changes that occur in what is called the beam helix, which projects along the cytoplasmic surface of each of the three Piezo1 arms. Flattening necessitates a kinking of the beam helix near S1330, whose role appears to complete a hydrogen bond in the kinked conformation. In support of this conclusion, when the serine is mutated to negate its hydrogen bonding capacity, mechanical gating is altered, requiring greater force to open the channel (Fig. 3G). Flattening of Piezo also causes a rigid body displacement of the beam helix toward the central axis of the channel, associated with the above-described conformational changes surrounding the pore (Fig. 3E and F).

### Inferences on the elasticity of Piezo1 in cell-derived membranes

If we look closely at the shape of Piezo1 in vesicles of different sizes we observe that in larger vesicles Piezo1 is less curved, that is, has a larger radius of curvature, R_p_ (Fig. 4A,B). In a previous study we documented the relationship between R_p_ and vesicle radius R_v_ by analyzing tomograms of individual Piezo1 vesicles made using reconstituted POPC:DOPS:cholesterol (8:1:1) lipids (Fig. 4C, black symbols)(*28*, *29*). In the present study using cell-derived membranes, after alignment of Piezo1 proteins in cryo-EM images to a consensus structure, proteins were re-extracted with a larger box size and 2D classification was performed. In this second step, 2D classification is driven by the vesicle shape, permitting us to separate channels into classes based on vesicle size (Fig. 4.A,B for outside-in Piezo vesicles and Fig. 4.C,D for outside-out Piezo vesicles). After making 3D reconstructions of Piezo1 from each individual vesicle size class, the Piezo1 radius of curvature for each class was approximated by fitting a circle to the middle of the channel’s density using side views, as shown for outside-in Piezo1 vesicles (Fig. 4B). Figure 4C shows R_p_ graphed as a function R_v_ taken from the four different vesicle sizes. The graph also shows analogous data from the tomographic analysis of outside-in Piezo1 channels in POPC:DOPS:cholesterol vesicles (Fig. 4C, black symbols). The data follow a very similar trend for vesicles comprising POPC:DOPS:cholesterol and cell-derived lipids. From this correspondence, we infer that the elastic properties of Piezo vesicles over the range of outside-in vesicle sizes analyzed here are similar in both the defined and cell-derived lipid compositions. Because the bending modulus of these two lipid membranes are likely similar(*35*), the bending modulus of the Piezo dome must also be similar in both.

The outside-out cell-derived Piezo1 vesicles approximate oblate spheroids with a flat Piezo1 channel (Figure 4D). Using the vesicle size classification approach described above, we find that outside-out Piezo1 channels adopt the same flat shape in vesicles with R_v_ = 27 nm and with R_v_ = 17 nm. From this observation we conclude that once Piezo1 reaches a certain degree of flattening, it no longer changes its shape when a greater flattening force is applied. To clarify this statement, recall that when Piezo1 is inserted into a vesicle in the outside-in orientation, the vesicle curvature squeezes Piezo1 into a conformation that is more highly curved than it would be in a planar membrane(*28*, *29*). In the outside-out orientation, because the vesicle membrane runs opposite to Piezo1’s intrinsic curvature, the vesicle membrane applies a flattening force onto Piezo1 that is more extreme in smaller vesicles. The force applied by an R_v_ = 27 nm vesicles is sufficient to maximally flatten Piezo1. We tried to image larger outside-out Piezo1 vesicles to estimate the elastic properties of Piezo1 as the completely flat shape is approached, but for technical reasons were unable to do this.

### Cell-derived lipids render a unique conformation of the flattened Piezo1 channel

We next compared the outside-out Piezo1 structure from cell-derived lipids in this study to a previously determined structure of Piezo1 reconstituted in an outside-out orientation in soy PC liposomes(*8*) (Fig. 5). Recall first that the functional data in Figure 1 show that Piezo1 is functional in cell-derived membranes, and not functional in soy PC membranes. Cryo-EM maps from similar sized vesicles show that Piezo1 remains curved (R_p_ about 100 nm) in PC membranes (Fig. 5B) and is completely flat (R_p_ –> ∞) in cell-derived membranes (Fig. 5A). It is as if something about the soy PC lipid environment prevents, or fails to facilitate, complete flattening of Piezo1.

**Figure 5.**
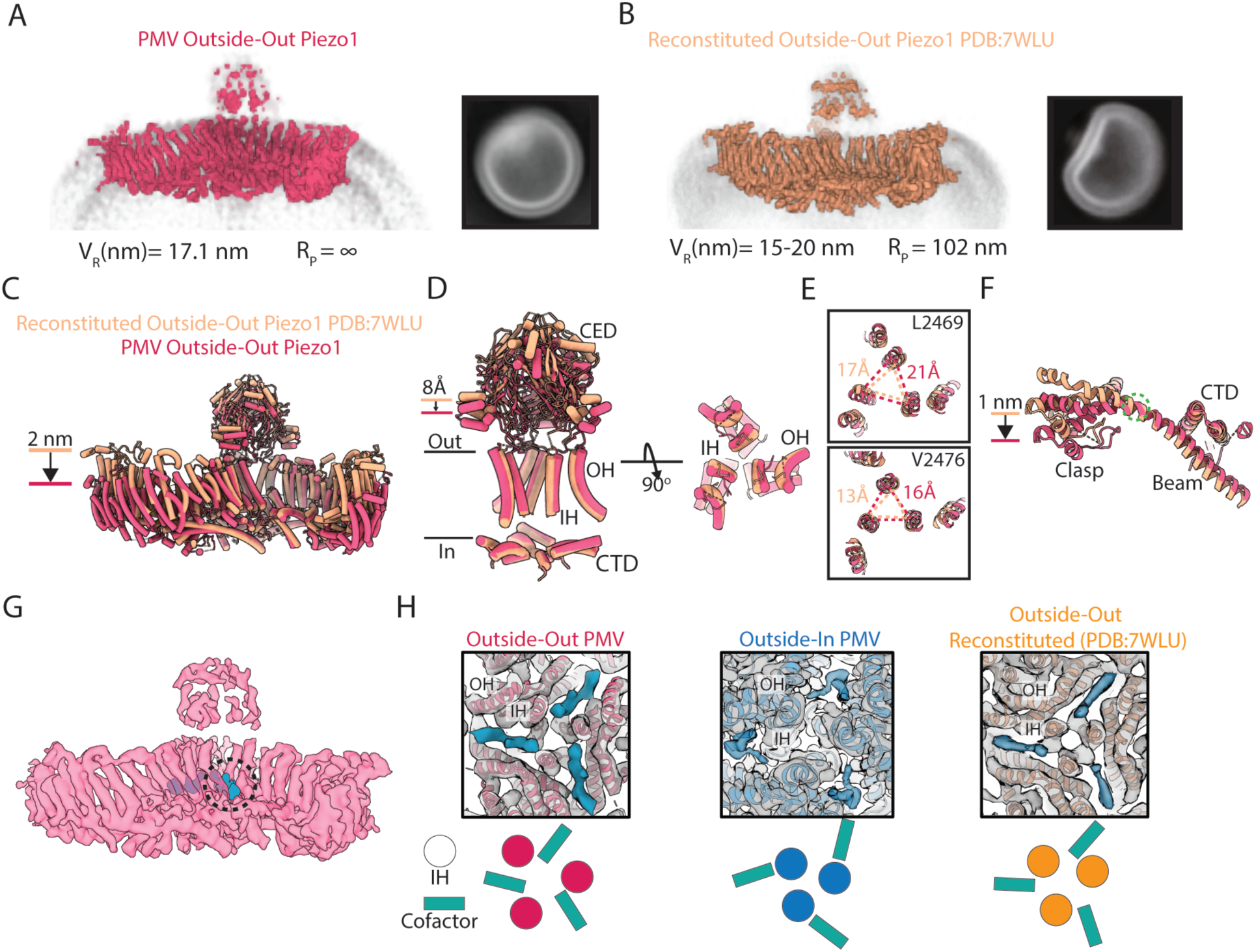
Comparison of the outside-out PMV and outside-out reconstituted Piezo1 structures. (A) (left) Cryo-EM map of outside-oriented Piezo1 PMV particles with a mean radius of 17.1 nm. The membrane is contoured in grey and the Piezo1 radius of curvature is labelled above. (right) 2D class image of these particles highlighting the flat edge of the vesicle where Piezo1 is. (B) (left) as in *A* but here of a Piezo1 structure determined by others in reconstituted liposomes^8^. (right) An image of a 2D class adapted from the reconstituted Piezo1 study^8^. (C) Aligned structures of the outside-out PMV (red) and reconstituted (orange) Piezo1 structures. The major displacement of the transmembrane arms is highlighted. (D) (left) Regions of Piezo1 involved in pore-gating are aligned for the outside-out PMV and reconstituted Piezo1 structures. The downward displacement of the CED as Piezo1 flattens is labelled. (right) The pore-lining helices are shown after a 90 ° rotation. (E) Two residue positions (L2469 and V2476) that form part of the hydrophobic constriction in the transmembrane pore region of Piezo1 are highlighted and the distances between their C*α* backbone residues are measured and annotated. (F) An isolated view of the Piezo1 beam helix, clasp and CTD region from the outside-out PMV and reconstituted structures are shown. Serine residue 1330, positioned proximal to a helical kink in the outside-out PMV Piezo1 structure, is highlighted by a dashed green circle. (G) Cryo-EM map of Piezo1 from the outside-out PMV dataset (red) with an unmodelled extra density highlighted in blue. (H) Top-down views of the Piezo1 transmembrane pore, highlighting the location of additional density relative to the IH and OH pore helices. Below each annotated structure is a schematic illustrating the relative position of the putative cofactor to the IH.

At the level of protein structure, complete flattening in cell-derived membranes compared to soy PC is associated with about a 2 nm displacement of the visible ends of the transmembrane arms (Fig. 5C). We summarize other conformational differences in Figure 5 C-F this way: to a first approximation, similar conformational changes occur in soy PC, but to a lesser extent. And our functional data suggest that completion of this transition may account for why Piezo1 is functional in cell derived membranes but not in soy PC or POPC:DOPS:cholesterol membranes. Together, these observations raise the question, what permits Piezo1 to complete the conformational transition in cell-derived membranes? Based on the data in Figure 4C and previous studies on the dependence of the membrane bending elastic modulus on lipid composition(*28*, *29*), we think the structural differences are unlikely due to differences in the elastic properties of the membranes.

One possible explanation for the structural and functional differences is that cell-derived membranes supply an essential ingredient. In our outside-out PMV maps we observe a prominent density that approaches the channel laterally from the inner leaflet of the bilayer, intercalating into the transmembrane pore between two adjacent IHs (Fig. 5G and H). The shape of the density is consistent with a lipid headgroup and two branched acyl chains, but the density is not well enough resolved to assign a specific lipid. An electrostatic representation of the Piezo1 surface in this region reveals an electropositive region that appears to cup one end of the density, consistent with a binding site for a phospholipid headgroup (S. fig 8). Indeed, several conserved lysine residues are near this putative lipid binding site (S.fig 8), including K2182-K2185, which were identified as forming a potential phosphoinositide interaction site by molecular dynamics studies(*36*). Mutation of all four of these lysines to alanine, asparagine or aspartate caused a rightward shift in the pressure-dependent activation threshold of Piezo1 in two separate studies(*36*, *37*), supporting a potential role of this region to Piezo1 gating. In structures of Piezo1 determined after reconstitution into liposomes, a ‘pore lipid’ has been built into density observed at this region, although its signal is weaker(*8*) (Fig. 5H). Here, the density resides just outside the pore, possibly reflecting a weaker binding interaction at this site (Fig. 5H). In our curved Piezo1 structure, we observe a similar density at this region, but here it is fully occluded from accessing the central pore by nearby hydrophobic amino acids, suggesting that pore expansion may be required for intercalation of the cofactor between the IHs.

## Discussion

Since the discovery of the Piezo channels, scientists have pursued an understanding of their force-dependent activation. In contrast to the prokaryotic mechanosensitive channels, which couple a large in-plane expansion associated with wide pore opening(*38–40*), Piezo1 channels activate at low tensions to form a cation-selective pore with a modest single channel conductance(*1*, *41*). Piezo1 also exhibits complex gating behaviors including voltage-dependent inactivation, which varies dramatically in different cell types(*1*, *32*, *42*, *43*). These properties likely enable the rich diversity of force-sensing processes Piezo1 is integral to. It is thus a worthwhile endeavor to understand how the Piezo channels are controlled by mechanical forces and other cues in living cells.

The first structures of Piezo1 determined in detergent, which revealed a highly curved, propeller-shaped trimeric channel, led to the membrane dome model(*6*). In this model, the deformation of the membrane bilayer by the Piezo1 transmembrane arms imbues the channel with its high-tension sensitivity, as flattening would be associated with a very large in-plane expansion of the membrane dome. This would have to be conformationally coupled to a more modest expansion of the channel’s pore, allowing the cation-selectivity of Piezo1 we observe. Further structural studies of Piezo1 reconstituted into liposomes revealed that it can flatten under force(*8*, *9*), and quantitative studies of the Piezo1 shape as a function of vesicle size estimated that in a planar bilayer, like a cell membrane under nominal tension, the Piezo dome adopts an intrinsic radius of curvature of about 40 nm. More recently MINFLUX microscopy studies on cell membranes corroborated these findings by showing a range of Piezo1 inter-blade distances consistent with a flatter conformation of Piezo1 than in detergent micelles(*44*).

Whilst these studies have demonstrated that Piezo1 can change its curvature(*8*, *9*, *44*), and even provided a quantitative relationship between Piezo1 shape and force(*28*, *29*), how mechanical sensing is transduced to open the pore and what properties give rise to the different gating behaviors of the channel in different physiological contexts has remained unclear. Central to our remaining ignorance is the fact that structural studies have been conducted under conditions that do not recapitulate force-dependent ion channel activity. Here, using single particle cryo-EM analysis of Piezo1 in cell-derived vesicles, where we show ion channel function is maintained, we describe conformational changes that can explain the activation of Piezo1 under force. Our most important observation is, in cell derived membranes Piezo1 can flatten to a degree, with pore-associated conformational changes, not seen previously in compositionally defined lipid membranes under otherwise similar conditions. These conformational changes are summarized in movie S1.

Recently, other studies have leveraged mutagenesis of Piezo1 to better understand the channel’s gating mechanism(*34*). Mutation of a serine residue to a glutamate (S2472E) within the transmembrane pore increased basal whole-cell poking currents of Piezo1 and slowed channel inactivation. One structure of Piezo1-S2472E in detergent was proposed to represent an intermediate conformation of channel opening. Based on alignments to our flattened structure we observe that the mutation has likely disrupted the conformation of the pore, uncoupling the force-dependency of Piezo1 from conformational changes of the gating apparatus (fig. S9). Whilst the pore-lining helices have expanded in the mutant structure, the conformation is distinct from what we observe in the flattened structure here and is not coupled to changes at the cytosolic gating apparatus (fig. S9).

What enables the fully-flattened Piezo1 channel conformation we observe in cell-derived membranes? Our analyses suggest that a cofactor present in cell membranes and absent in previous reconstituted studies, is necessary for complete flattening and the associated conformational changes to the Piezo1 gating apparatus we observe. The extra density we observe in the flattened structure is a good candidate for such a cofactor, and its properties are most consistent with a lipid molecule. The absence of this lipid in previous structures determined is consistent with the absence of mechanical activation of Piezo1 in identical conditions, as observed by patch-clamp recordings even under large applied pressures. Both ceramide and PIP_2_ have been shown to be functionally important for Piezo1 activation(*42*, *45*) and molecular dynamics simulations implicate an abundance of interaction sites to the channel(*46*, *47*). More work will be needed to identify and validate the role of this cofactor. The idea that Piezo1 is primarily mechanosensitive but requires a cofactor for activation is very reminiscent of some other ion channels that exhibit multiple modes of gating. The Kv7.1 K^+^ channel is a good example(*48*, *49*). It’s opening is voltage-dependent but requires PIP_2_ to open. Membrane depolarization alone is insufficient.

A simple schematic for a multi-modal gating of Piezo1 is shown in Figure 6. To achieve the activated conformation both mechanical force and the cofactor are required. In the depiction, the two events – mechanical force and the cofactor binding – occur sequentially, but this point needs further study. The requirement of two stimuli might rationalize the variable, environment-dependent properties of Pizeo1 function, such as channel inactivation, observed in different cell types and under different conditions(*1*, *32*, *42*, *43*). There is evidence for both lipid-mediated(*42*, *50*) and protein-mediated modulation of Piezo1 function(*43*, *51*), with some described mechanisms being cell-type specific(*42*, *43*). Most recently, the transcriptional regulator, MyoD (myoblast determination)-family inhibitor domain-containing (MDFIC), which abrogates channel inactivation, was shown to bind at the same regulatory site postulated in the present study^43^(fig. S10). Although the physical basis for Piezo1 channel inactivation is still unknown, it is tempting to hypothesize that this binding site serves as a nexus for cues additional to mechanical force that regulate Piezo1 function, enabling the rich diversity of force-sensing processes Piezo1 has been ascribed to.

**Figure 6.**
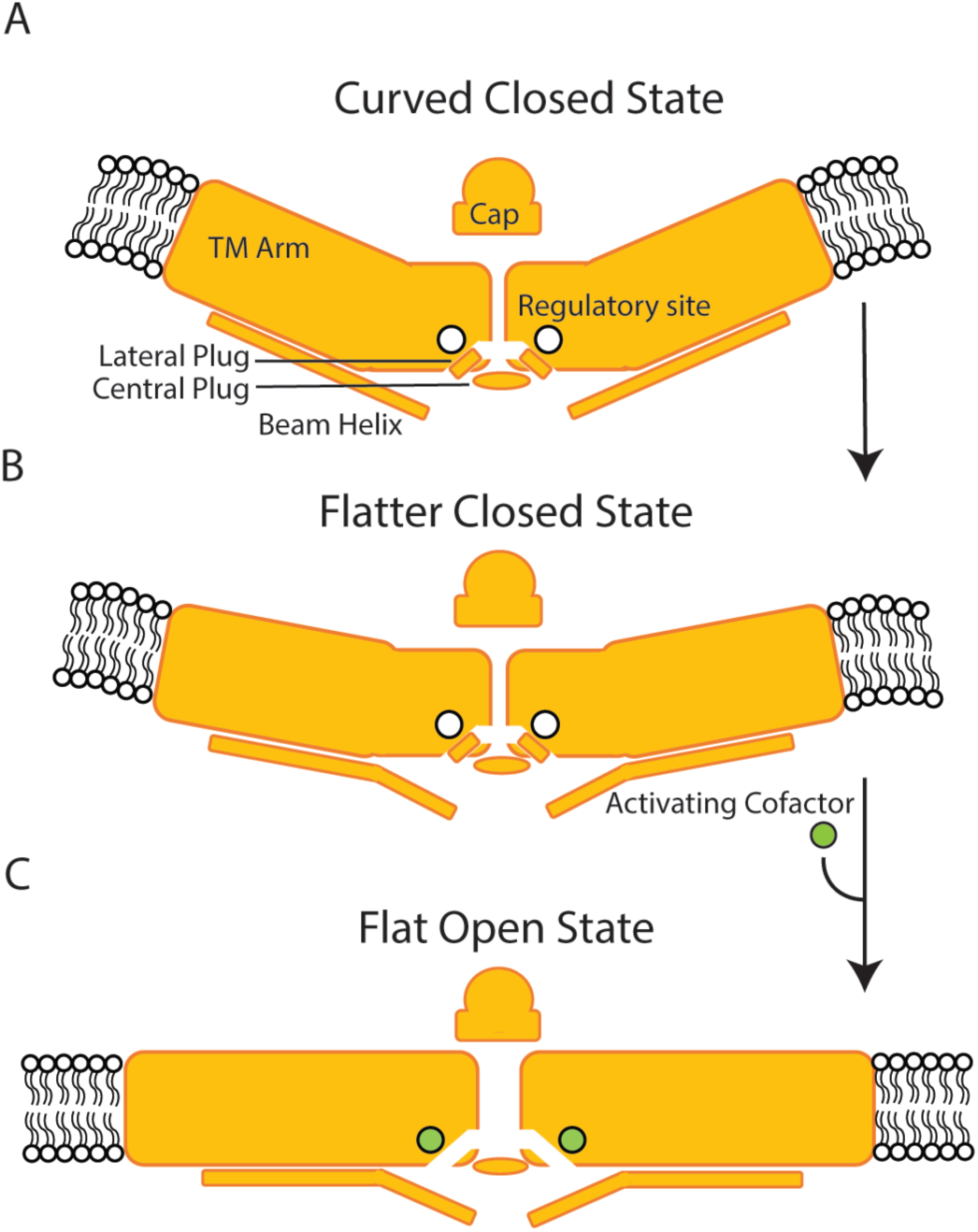
Proposed mechanism for multi-modal gating of Piezo1. (A) Under resting conditions in an approximately planar cell membrane, Piezo1 adopts a slightly curved conformation with an intrinsic radius of curvature of approximately 42 nm. (B) With increasing membrane tension, the transmembrane arms of Piezo1 begin to flatten, supported by the beam helix. Without occupancy of the regulatory site the channel pore remains closed. (C) At some given mechanical force, in combination with binding of the activating cofactor at the regulatory site, Piezo1 flattens fully into an activated conformation, associated with displacement of the lateral plug, enabling ion conduction.

## Acknowledgements

We thank all members of the MacKinnon lab for their support and feedback. We would like to thank Mark Ebrahim, Johanna Sotiris and Honkit Ng at the Evelyn Gruss Lipper Cryo-EM Resource Center at Rockefeller University for assistance with cryo-EM grid screening. We thank Rui Yan and staff members at the HHMI Janelia Cryo-EM Facility for their assistance with cryo-EM microscope operation and data collection. RM is an investigator at the Howard Hughes Medical Institute.

## Author contributions

GV and RM conceptualized and designed the project. GV performed all experiments. GV and RM analyzed the data. GV and RM wrote and edited the manuscript.

## Supplementary Materials

Materials and Mthods

Figs. S1 to S10

Table S1

References 52 to 56

Movie S1

## Materials and Methods

### Cell culture and transfection

PIEZO1^-/-^ HEK293T cells were generated previously by crispr-CAS9 technology in the lab^18^ and were cultured in DMEM medium (Gibco) supplemented with 10% fetal bovine serum. *Spodoptera frugiperda* Sf9 cells (ATCC) were cultured in Sf-900 II SFM medium supplemented with 100 U/ml penicillin and 100 U/ml streptomycin at 27 °C. Expi293F cells (Thermofisher) were cultured in Expi293 expression medium (Thermofisher)

### Protein Expression

Piezo1 was heterologously expressed in Expi293F cells using the BacMam method as previously described^2,3^. Bacmids were generated by transforming the mPiezo1-ALFA_2420_-HRV3C-GFP construct into *Escherichia coli* DH10Bac cells. Baculoviruses were then produced by transfecting Sf9 cell with the bacmid using Cellfectin II (Invitrogen). After two rounds of amplification, baculoviruses were used for cell transduction. Expi293F cell suspension cultures were grown at 37 °C to a density of ∼3 x 10^6^ cells/mL and infected with 10% (v:v) of the baculovirus. After 16 h, 10 mM sodium butyrate was supplemented, and the temperature was shifted to 30 °C. Cells were harvested ∼48 h later. For protein purification and reconstitution, cell pellets were flash-frozen in liquid nitrogen and used later. For NEM-induced membrane vesicle (PMV) isolation, cells were used immediately without freezing.

### Purification of Piezo1 in detergent

Cell pellet from 2L of Expi293F cells expressing mPiezo1-ALFA_2420_-HRV3C-GFP were resuspended in 200ml of buffer containing 150 mM NaCl, 20 mM Tris pH8 (TBS) and a protease inhibitor mix containing 0.1 µg/mL pepstatin A, 1 µg/mL leupeptin, 1 µg/mL aprotinin, 1 mM benzamidine, 0.1 mg/ml 4-(2-aminoethyl) benzenesulfonyl fluoride hydrochloride (AEBSF), and 1 mM phenylmethylsulfonyl fluoride hydrochloride (PMSF). Resuspended cells on ice were lysed by sonication with a probe sonication (1/2” tip with the Branson102-C converter) at 40% power for 6 x 30 s pulses, with 30 s delays between. After a 5000G spin for 15 minutes, the supernatant was ultracentrifuged at 100,000 g at 4 °C for 40 mins. The resultant membrane pellet was resuspended in TBS (5ml/ pellet) and dounce homogenized for ∼ 20 strokes. The membrane suspension was solubilized in TBS buffer containing 2% C12E10 and protease inhibitors for 90 mins at 4 °C. The solubilized suspension was then ultracentrifuged at 100,000 g at 4 °C for 40 mins and the resultant supernatant was incubated with 1ml ALFA Selector CE resin (NanoTag), preequilibrated with TBS buffer containing 0.025% C12E10 (SEC buffer), at 4°C overnight.

The following day, the ALFA selector CE resin was batch-washed twice with ∼20 mL SEC buffer. The resin was then loaded onto a gravity column and washed with another 15 ml SEC buffer. Protein was then eluted in 10 mL SEC buffer containing 0.2 mM ALFA peptide (NanoTag) by incubation at 4 °C for 2 hours. Eluted protein was concentrated to ∼500 µL using an Amicon 4 mL concentrator (molecular weight cutoff of 30 kDa). After a 5000 G spin for 5 minutes at 4°C to pellet aggregate, the supernatant was further purified on a Superose-6 size exclusion column (10/300 GL) in SEC buffer at 4°C. The peak fractions corresponding to the trimeric mPiezo1 were concentrated to ∼ 2 mg/mL using a 2 mL Amicon concentrator (molecular weight cutoff of 30 kDa) to be used for reconstitution into liposomes.

### Reconstitution of Piezo1 into liposomes

Soy PC (Avanti) lipids were resuspended in chloroform at a concentration of 20 mg/mL and dried to a thin film under argon. The lipid film was further dried overnight in a room-temperature vacuum desiccator and then resuspended at a concentration of 10mg/mL by mixing with TBS buffer. Small unilamellar vesicles were formed by bath sonication at RT until the solution was mostly transparent and the suspension was solubilized by addition of 2% n-decyl-ß-D-maltoside (DM) (Avanti) for 30 min at room-temperature. Solubilized lipids were mixed with concentrated Piezo1 at protein:lipid ratios of 1:50 and 1:500 for 1 hour at 4 °C. Detergent was removed using adsorbent Bio-Beads SM-2 Resin (Bio-Rad) by adding 20 mg of a 50% (wt/vol) Bio-Beads slurry in TBS buffer and rotating at 4 °C overnight. The proteoliposomes were then concentrated ∼ 5x in a 500 µL Amicon spin concentrator (molecular weight cutoff of 100 kDa). The concentrated proteoliposomes were then flash-frozen at liquid nitrogen to use later.

### Electrophysiology

For excised outside-out patch clamp recordings, 3 µg of plasmid was transfected into HEK293 cells at ∼50-60% confluency using FuGENE HD transfection reagent following the manufacturer’s instructions (Promega). Recordings were made 24h to 48h posttransfection.

For patch clamp recordings of giant plasma membrane vesicles (GPMVs), cells were first transfected with plasmid as described above. Following expression, DMEM media was removed from the adherent HEK293 cells which were then washed twice with GPMV buffer: 150 mM NaCl, 5 mM KCl, 20 mM Na-HEPES and 2 mM CaCl_2_, pH7.4 before then being incubated with GPMV buffer + 7.5 mM N-ethylmaleimide (NEM) for 2 hours at 37 °C. The solution was then aspirated using a 1000 µL pipette with the tip cut off and allowed to sit at RT for 10 minutes to pellet detached cells and free GPMVs. 20 µL of GPMVs, aspirated from just above the cell pellet, was then added to bath solution in a glass-bottomed dish.

For patch clamp recordings of reconstituted Piezo1, giant uinlamellar vesicles were formed from Piezo1-containing proteoliposomes. 10 µL of proteoliposome solution was pipetted into ∼6 spots on a glass-bottomed dish and dried for at least 6 hours at RT in a vacuum desiccator. 50 µL of rehydration buffer: 140 mM NaCl and 20 mM Na-HEPES, pH 7.4 was then added to the dried down spots and the dish was placed into a 150 x 25 mm dish containing wet Kimwipes and allowed to rehydrate overnight at 4 °C. The following day, bath solution was added and after 30 minutes, the sample was ready for patch clamping.

For all electrophysiology experiments, bath solution and pipette solution consisted of 140 mM NaCl and 20 mM Na-HEPES at pH 7.4. Imaging was performed on a Nikon eclipse DIC microscope at x40 magnification. Pipettes of borosilicate glass (Sutter Instruments; BF150-86-10) were pulled to ∼2-5 MΩ resistance with a micropipette puller (Sutter Instruments; P-97) and polished with a microforge (Narishige; MF-83). Recordings were obtained with an Axopatch 200B amplifier (Molecular Devices), filtered at 1 kHz and digitized at 10 kHz (Digidata 1440A; Molecular Devices). A high-speed pressure clamp (ALA scientific) was used to form gigaseals and reversibly apply pressures to activate Piezo1 channels. For excised recordings from GPMVs and GUVs, if the vesicle remained attached after gigaseal formation and pulling away of the pipette, the vesicle would be lifted briefly to the air-water interface to remove the attached vesicle but leave an intact membrane patch inside the pipette.

### Piezo1 PMV preparation and purification

After expression of Piezo1 by baculovirus transduction as described above, 4L of live cells was harvested by centrifugation at 3,000 g for 10min. The cell pellet was resuspended in 400 mL GPMV buffer containing 140 mM NaCl, 5 mM KCl, 10 mM Na-Hepes and 2 mM CaCl_2_, and 7.5 mM NEM at pH 7.4. The cell suspension was incubated in baffled flasks at 37 °C, shaking at 130 rpm for 2 h. After incubation, the flasks were shook vigorously by hand for ∼20s, to release more vesicles, before spinning at 3,000g at 4°C for 10 min to pellet cells and GPMVs.

The supernatant was then supplemented with a protease inhibitor mix containing 0.1 µg/mL pepstatin A, 1 µg/mL leupeptin, 1 µg/mL aprotinin, 1 mM benzamidine, 0.1 mg/ml AEBSF, and 1 mM PMSF and 10% glycerol before sonication with a probe sonicator at 40% power for 30 sec pulses, 6 times with 30 sec breaks in between. The vesicles were spun at 3,000g for 10 mins to pellet aggregate before ultracentrifugation at 100,000 g at 4 °C for 30 min in a Ti70 rotor. The membrane pellet from every ∼25 mL sample was triturated in ∼1mL GPMV buffer supplemented with 10% glycerol. The resuspended vesicles were then sonicated in a bath sonicator (Branson M1800) for ∼30 s. A centrifugation step at 3,500 g for 10 min was then performed to pellet remaining aggregates. The following affinity purification steps then depended on whether we were isolating inside-out or outside-out Piezo1 PMVs.

For inside-out PMVs the vesicles were incubated with 2 mL GFP-Trap Magnetic Particles M-270 resin (Proteintech), preequilibrated with GPMV buffer, for 2 h at RT. The resin was then separated from flowthrough solution using a DynaMag-2 magnet and washed with 10 mL total GPMV buffer three times. The bound vesicles were then eluted by incubating with HRV-3C protease at a final concentration of ∼0.014 mg/mL at RT for 2 h. The eluted vesicles were concentrated in an Amicon 0.5 mL concentrator (molecular weight cutoff of 100 kDa) to an OD_280_ of ∼2-4.

For outside-out PMVs the vesicles were incubated with 2 mL ALFA Selector PE resin (NanoTag) preequilibrated with GPMV buffer at 4 °C overnight. The following day, the ALFA selector PE resin with bound vesicles was first batch-washed twice with ∼20 mL GPMV buffer and spun at 1,000 g at 4 °C for 1 min to collect resin, followed by a second wash with 20 mL GPMV buffer. The resin was then loaded onto gravity column and washed further with 20 mL GPMV buffer. The vesicles were eluted with 10 mL GPMV buffer supplemented with 0.2 mM ALFA peptide by incubating at RT for 1 h. Due to the persistent occurrence of inside-out Piezo1 PMVs and Piezo1-containing broken membrane “fragments” another purification step was included to sequester these contaminants. Eluted vesicles were incubated with 2 mL GFP-Trap Magnetic Particles M-270 resin (Proteintech), preequilibrated with GPMV buffer, for 2 h at RT. The supernatant was separated from the resin using a Dynamag-2 magnet and the flow-through, containing mostly outside-out Piezo1 PMVs, was concentrated in an Amicon 0.5 mL concentrator (molecular weight cutoff of 100 kDa) to an OD_280_ of ∼2-4.

### Grid preparation and Data Collection

Quantfoil R1.2/1.3 400 mesh holey carbon grids were glow-discharged for 22 s. Then, 3.5 µL concentrated membrane vesicles were applied to freshly glow-discharged grids and left for 5-8 min at 22 °C. The grids were blotted manually with a piece of filter paper before another 3.5 µL of sample was applied. The grids were blotted using a Vitrobot Mark IV with a blot force of 0 and blot time of 3 s after 20 s of incubation. The grids were flash-frozen in liquid ethane and stored in liquid nitrogen until data collection.

Both the inside-out and outside-out Piezo1 PMV datasets were collected on a 300 keV Titan Krios transmission electron microscope equipped with a cold-field emission gun and an energy filter (slid with 6 eV). For the inside-out PMV dataset a total of 19144 movies were recorded by a Falcon 4i camera with a physical pixel size of 0.94 Å and a target efocus value of -1.5 ∼ -2.5 µm. The movies have 2142 internal frames and a total dose of 60 e^-^/ Å^2^.For the outside-out PMV dataset, a total of 16060 movies were recorded by a Falcon 4i camera with a physical pixel size of 0.94 Å and a target efocus value of -1.5 ∼ -2.5 µm. The movies have 1953 internal frames and a total dose of 60 e^-^/ Å^2^.

### Cryo-EM Data processing

The data-processing workflow is detailed in figures S2 and S4. Data processing was carried out using cryoSPARC v4 (ref) and RELION 4.0 (ref). The movies were gain-normalized and motion corrected in cryosparc. Contrast transfer function parameters were estimated from the motion-corrected micrographs using the Patch CTF estimation tool. All subsequent processing was performed on motion-corrected micrographs with dose weighting.

For the outside-in PMV dataset (fig. S2), ∼500 particles were first manually picked, separately selecting sideviews and top views of Piezo1. These sets of particles were then used to train a TOPAZ picking model (ref), which was subsequently used to pick particles on approximately a fifth of the dataset. Good 2D classes were selected and used to retrain a TOPAZ picking model and this was done iteratively to improve the picking model. A combined total of 529,310 particles were picked from the side and top views. The best side view 2D classes were picked and an initial map generate by ab initio reconstruction. This map was refined by homogenous refinement in C3 and used as a reference for heterogeneous refinement to sort for the well aligned particles which were then refined in C3. Next, a seeded heterorefinement method was used to enrich for good particles (ref) and include top-views. In brief, particles selected from good 2D classes of top-views and sideviews were split into subgroups, combined with the well-aligned particles seed particles selected from previous steps and hetereogeneous refinement was performed. For each round, the good classes were selected, the redundant particles removed and refinement performed. To improve the density of the CED, the CED and central pore unit were masked out and locally refined in C3. To improve the density of the transmembrane arms, the particles were symmetry expanded in C3 and one arm with the CED were masked out before local refinement was performed in C1. These focused maps were then combined in Phenix to generate the final reconstruction.

For the outside-out PMV dataset (fig. S4), ∼500 particles were first manually picked and used to train a TOPAZ picking model which was used to re-pick particles and iterative rounds of this were performed to achieve the best TOPAZ picking model. From these particles, 2D classification was performed. Despite extensive efforts, we were unable to get good 2D classes of top-down views. From the best ∼100,000 particles, hetereogenous refinement was performed using a 60 Å low-pass filtered map of a previously determined structure of outside-out oriented Piezo1 reconstituted into liposomes (EMD-32593)(ref). Multiple rounds of heterorefinement were performed this way. The best-aligned particles and map were used for a seeded heterorefinement strategy, as discussed above, enriching the number of well-aligned particles approximately six-fold. After combining the good particles and removing redundante ones, C3 refinement was performed and these particles were expanded by C3 symmetry. One transmembrane arm and CED were masked out RELION (ref) 3D classification was used without alignment in C1 to separate out the bad particles. The best particles then went through another round of 3D classification, this time with local angular searches and the best particles were then imported into cryosparc, the duplicates removed and local refinement in C3 used to achieve the final map.

### Model Building and Refinement

A structural model for the inside-out Piezo1 PMV was built by docking in the structure of Piezo1 in detergent micelle (PDB ID:6B3R)^6^ and adjusted using the ISOLDE plugin^52^ in Chimera X^53^ or Coot^54^ followed by real-space refinement in Phenix^55^. The outside-out Piezo1 PMV structural model was built using a starting structure of Piezo1 reconstituted into liposomes (PDB ID: 7WLU)^8^, that was fit into the map by rigid-body refinement in ISOLDE and Phenix. The quality of the final models was evaluated using the MolProbity plugin^56^ in Phenix. Graphical representations of models and cryo-EM maps were prepared using ChimeraX. Map and structure statistics for the two structures are given in the supplementary materials, Table S1.

---

**Fig. S1.**
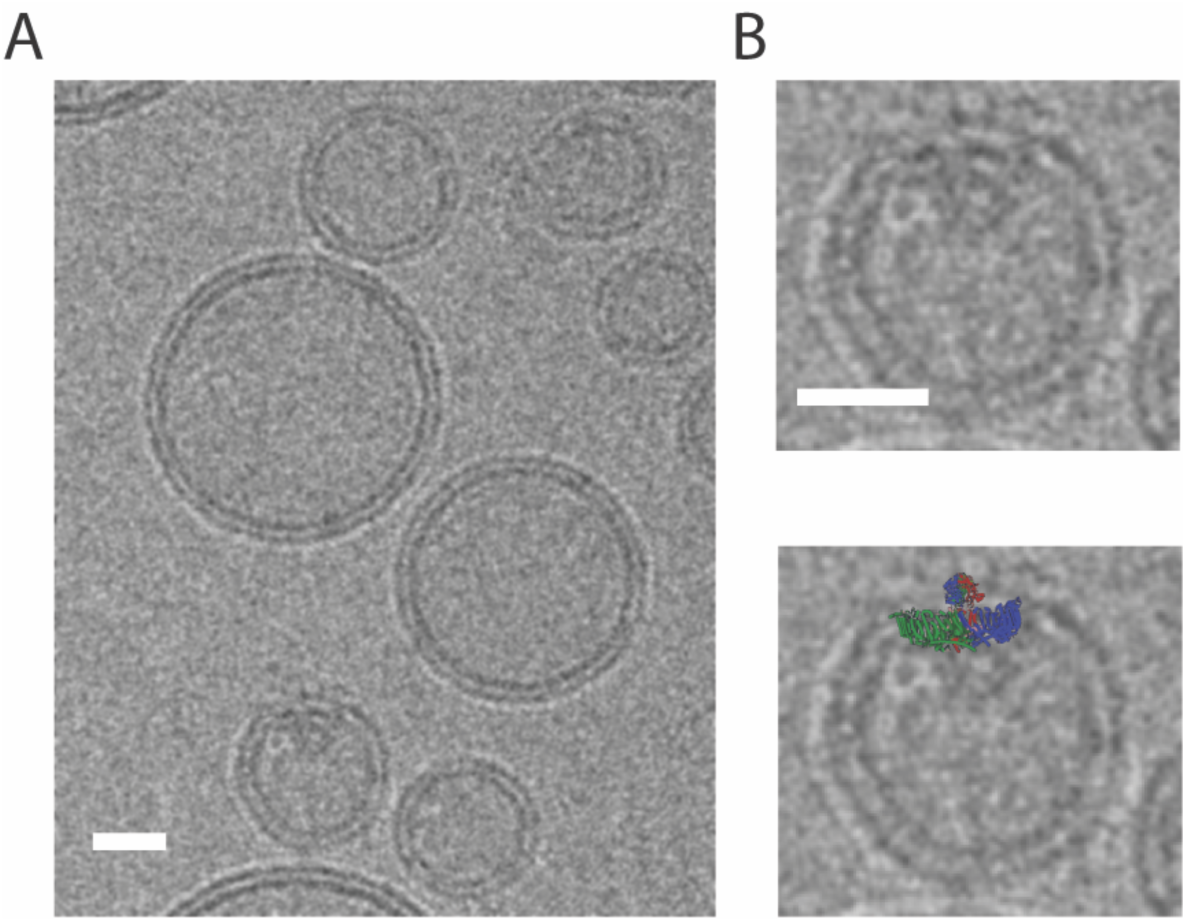
Reconstitution of purified Piezo1 into soy PC liposomes. (A) Raw micrograph image from electron microscopy of Piezo1 reconstituted into soy PC liposomes. (B) Close-up of a liposome containing a Piezo1 channel with a structure of Piezo1 overlaid for clarification. Scale bar = 20 nm.

**Fig. S2.**
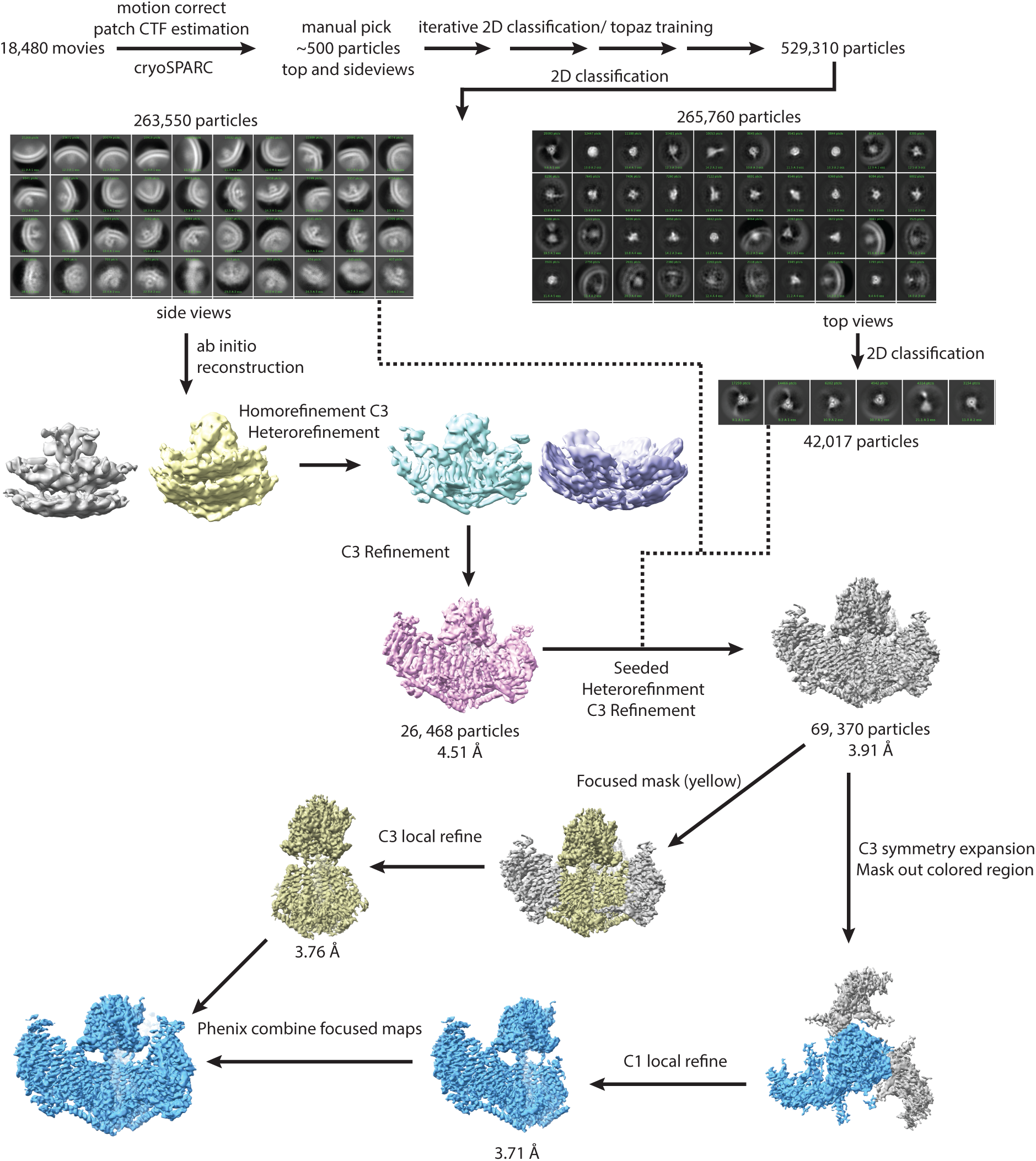
Single particle cryo-EM processing workflow for the outside-in Piezo1 PMV dataset.

**Fig. S3.**
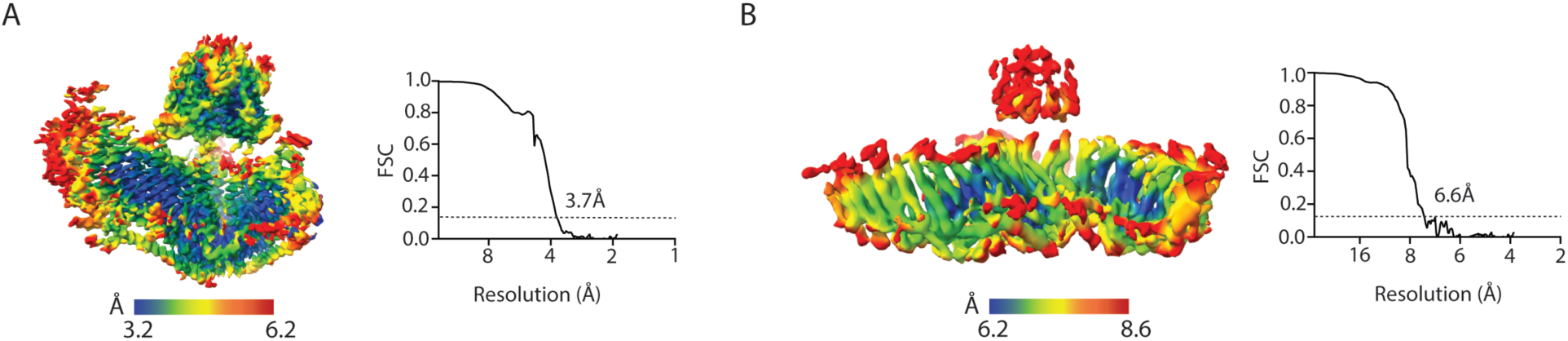
Local resolution maps and FSC curves. The local resolution maps and FSC curvs are generated by CryoSPARC. **(A)** Outside-in Piezo1 PMV. **(B)** Outside-out Piezo1 PMV.

**Fig. S4.**
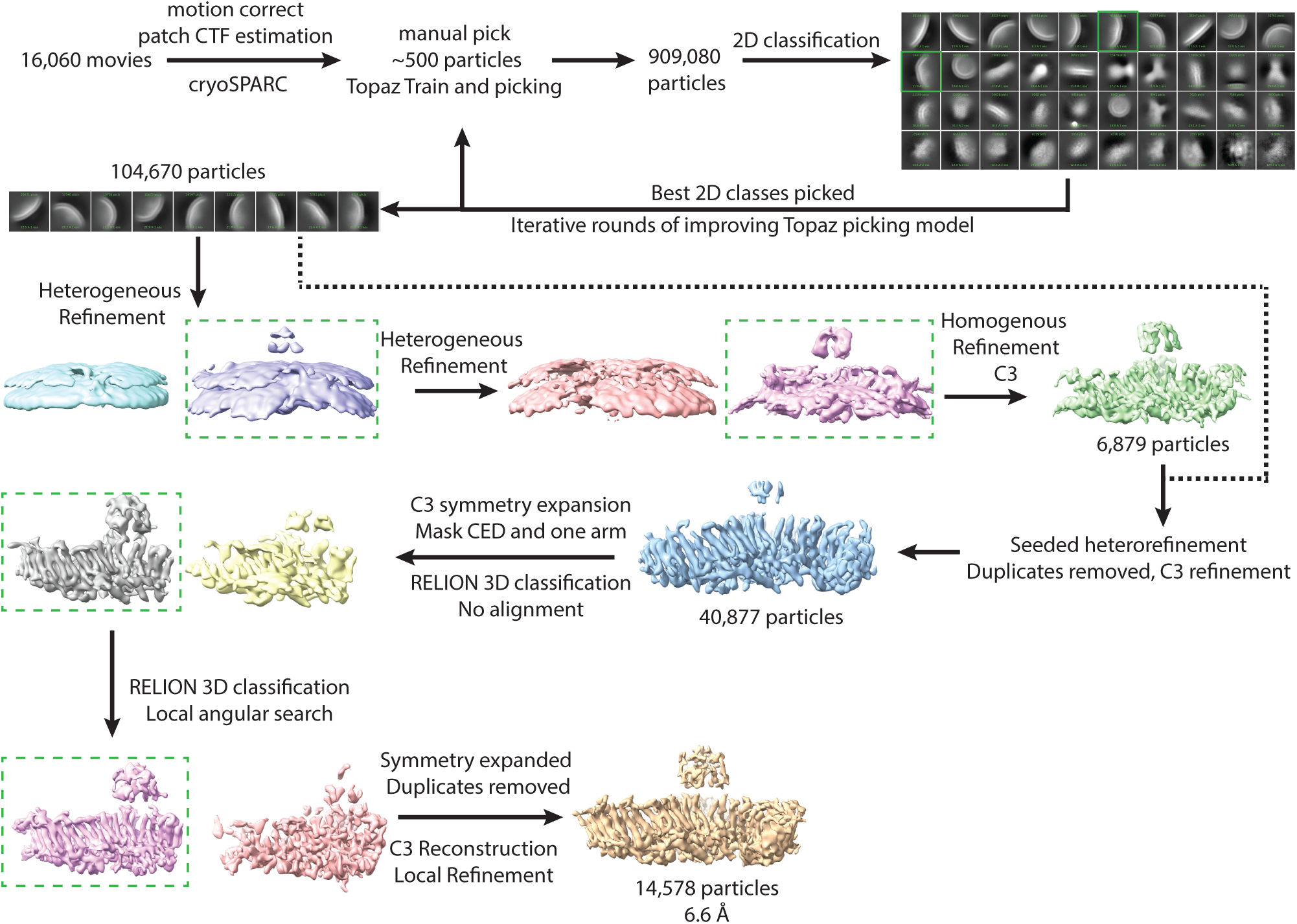
Single particle cryo-EM processing workflow for the outside-out Piezo1 PMV dataset.

**Fig. S5.**
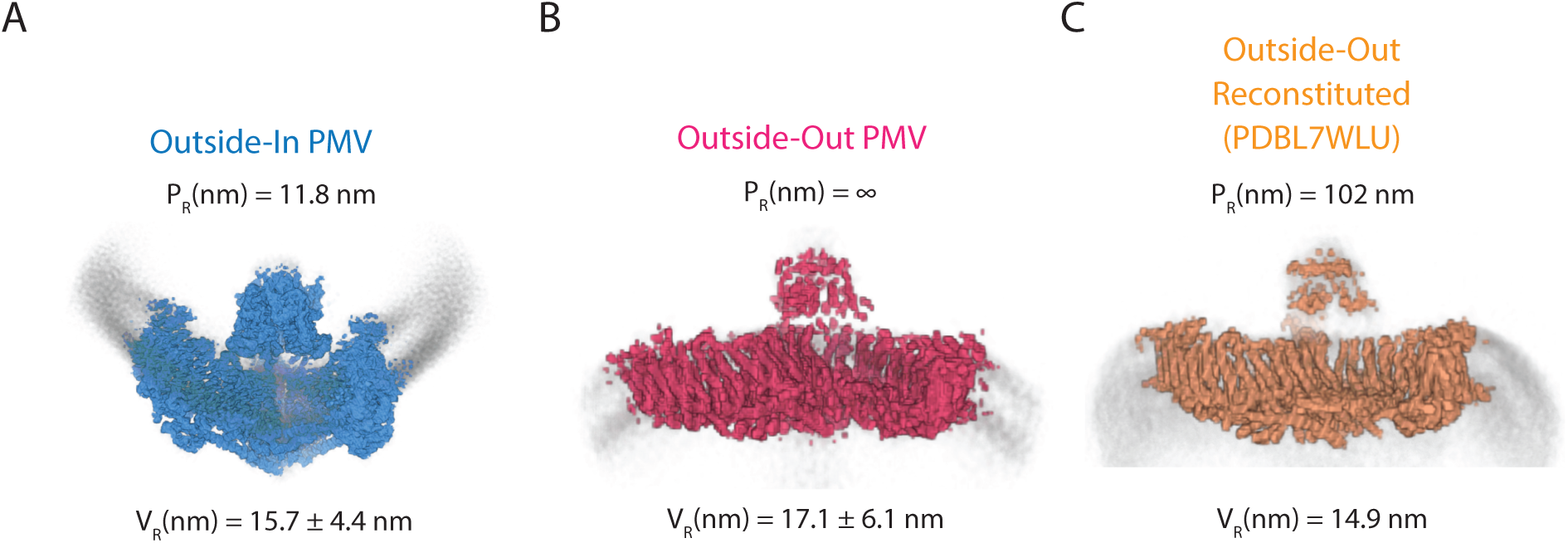
Cryo-EM maps of Piezo1 in vesicles. **(A)** Cryo-EM map of outside-in oriented Piezo1 in PMVs with the membrane contoured in grey. The Piezo1 radius of curvature and the mean radius of the vesicles contributing to this structure are annotated. **(B)** as in *A* for the outside-out Piezo1 PMV structure. **(C)** As in *A* for a structure determined by others of Piezo1 reconstituted in the outside-out orientation in liposomes^8^.

**Fig. S6.**
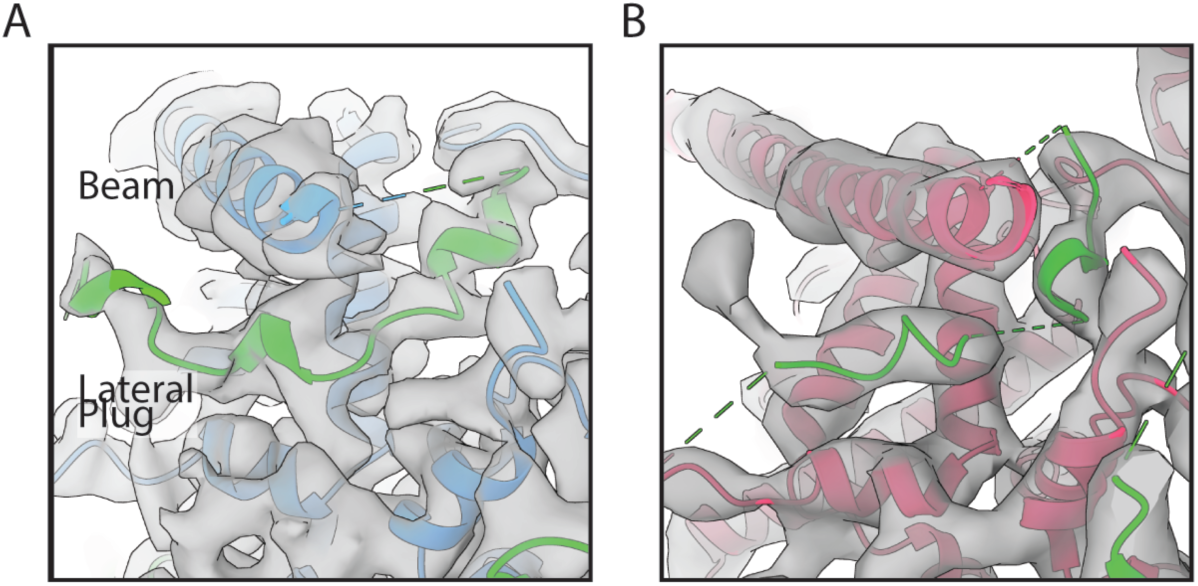
Cryo-EM map of the Lateral Plug. Close-up view of the lateral plug structure (green) and cryo-EM map (gray) for the outside-in (A) and outside-out (B) PMV Piezo1 structures.

**Fig. S7.**
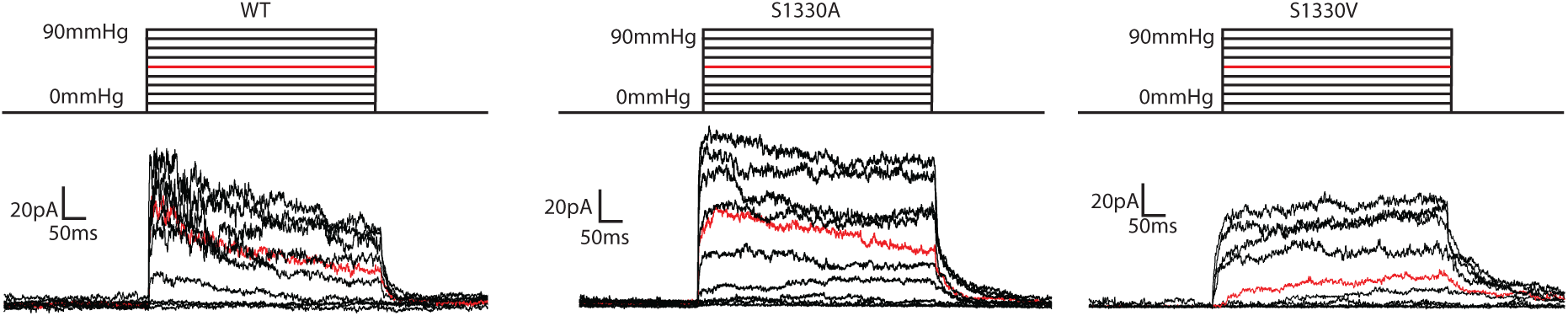
Representative pressure-activated currents of WT Piezo1 compared to two mutations made at a serine residue in the beam helix. Outside-out patch clamp recordings were made from Piezo1 knockout HEK293 cells overexpressing either WT Piezo1 or one of the mutants S1330A or S1330V. Currents were held at +60 mV and stepwise 500ms pressure pulses from 0 to -90mmHg in -10mmHg increments were made. The red trace depicts the current elicited at -50 mmHg.

**Fig. S8.**
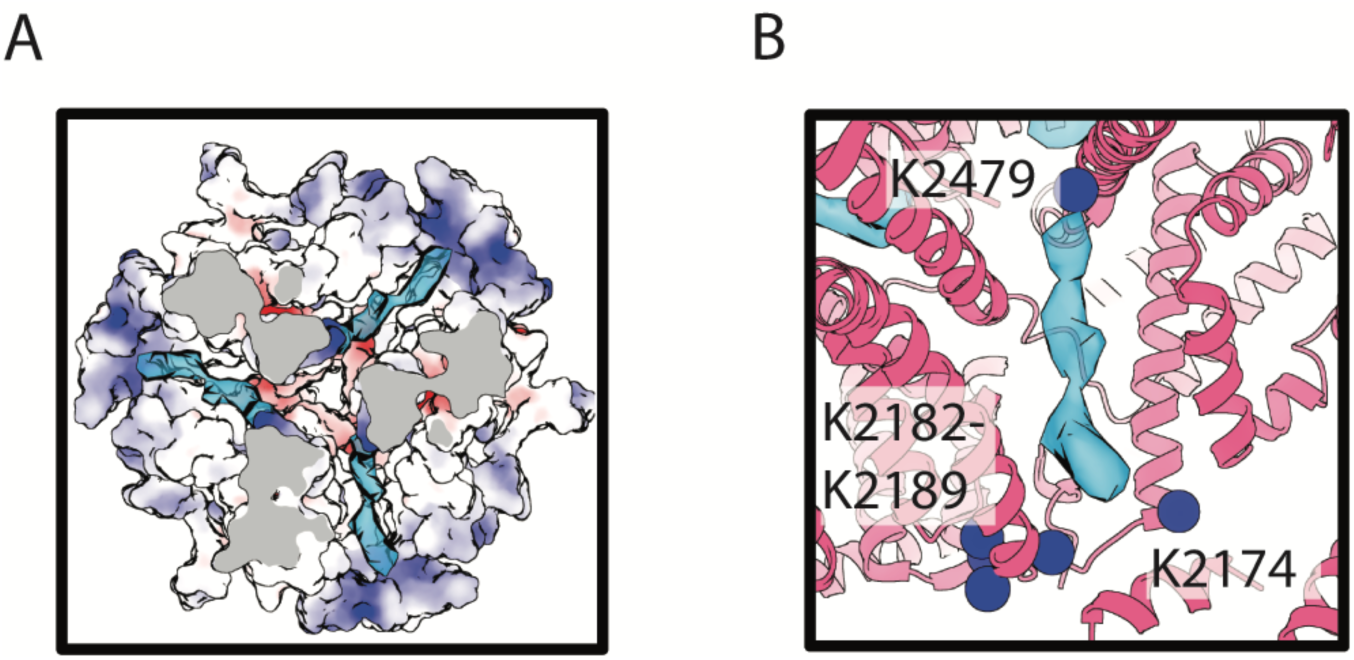
Focus on the Piezo1 regulatory binding site. (A) Top-own view of the outside-out PMV Piezo1 transmembrane pore from at the regulatory binding site, depicting the electrostatic surface, where blue is electropositive, white is neutral and red is electronegative. The cryo-EM map of the putative cofactor is included. (B) A close-up view of the binding site, depicting the positions of proximal lysine residues as blue spheres relative to the cryo-EM map of the putative cofactor.

**Fig. S9.**
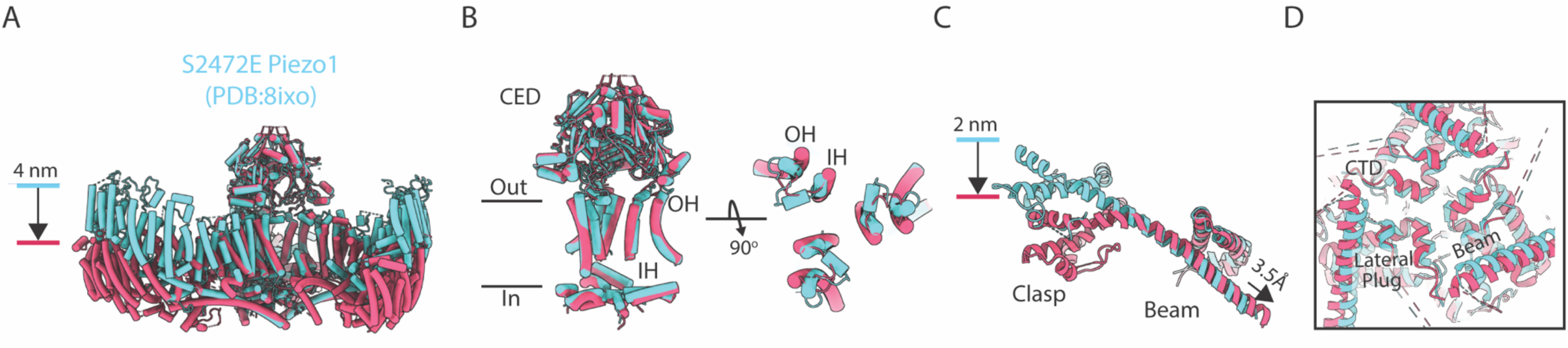
Comparison of outside-out PMV Piezo1 and a Piezo1 mutant, S2472E. (A) Aligned structures of outside-out PMV (red) and Piezo1 mutant S2472E (cyan). The major displacement of the transmembrane arms is highlighted. (B) Regions of Piezo1 involved in pore-gating are aligned from a side-view (left) and top-down view with a close-up of the pore-lining helices (right). (C) An isolated view of the Piezo1 beam helix, clasp and CTD region of the two structures. (D) A close-up view of the beam, CTD and lateral plug that has been proposed to form the cytosolic gate for ion conduction in Piezo1, is shown. The sliding of the beam helix is associated with conformational changes at this region in the outside-out Piezo1 structure and not the mutant structure.

**Fig. S10.**
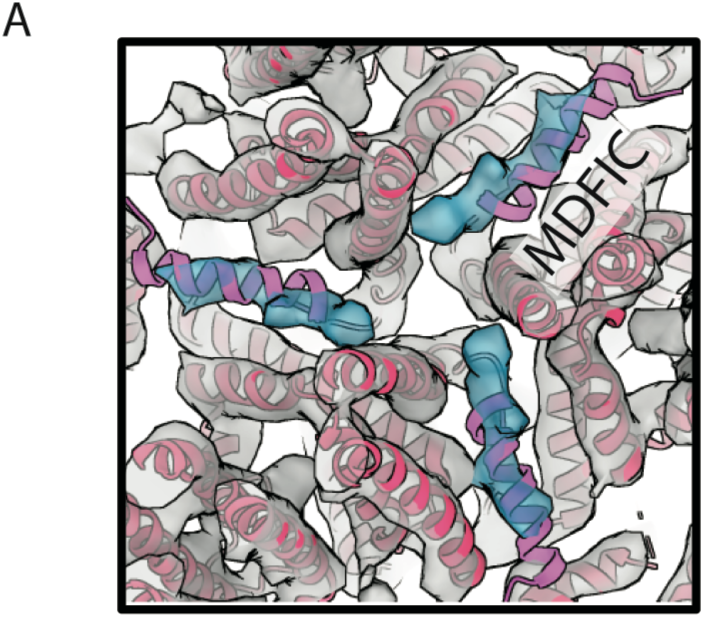
An auxiliary regulator of Piezo1 binds to the regulatory site. A top-down view of the binding site in the outside-out PMV Piezo1 structure (red), the corresponding cryo-EM map (grey) and the bound putative lipid density (blue) with the C-terminal helix of MDFIC from a Piezo1-MDFIC complex structure (PDB:8IMZ) aligned and shown in pink.

**Table S1.**
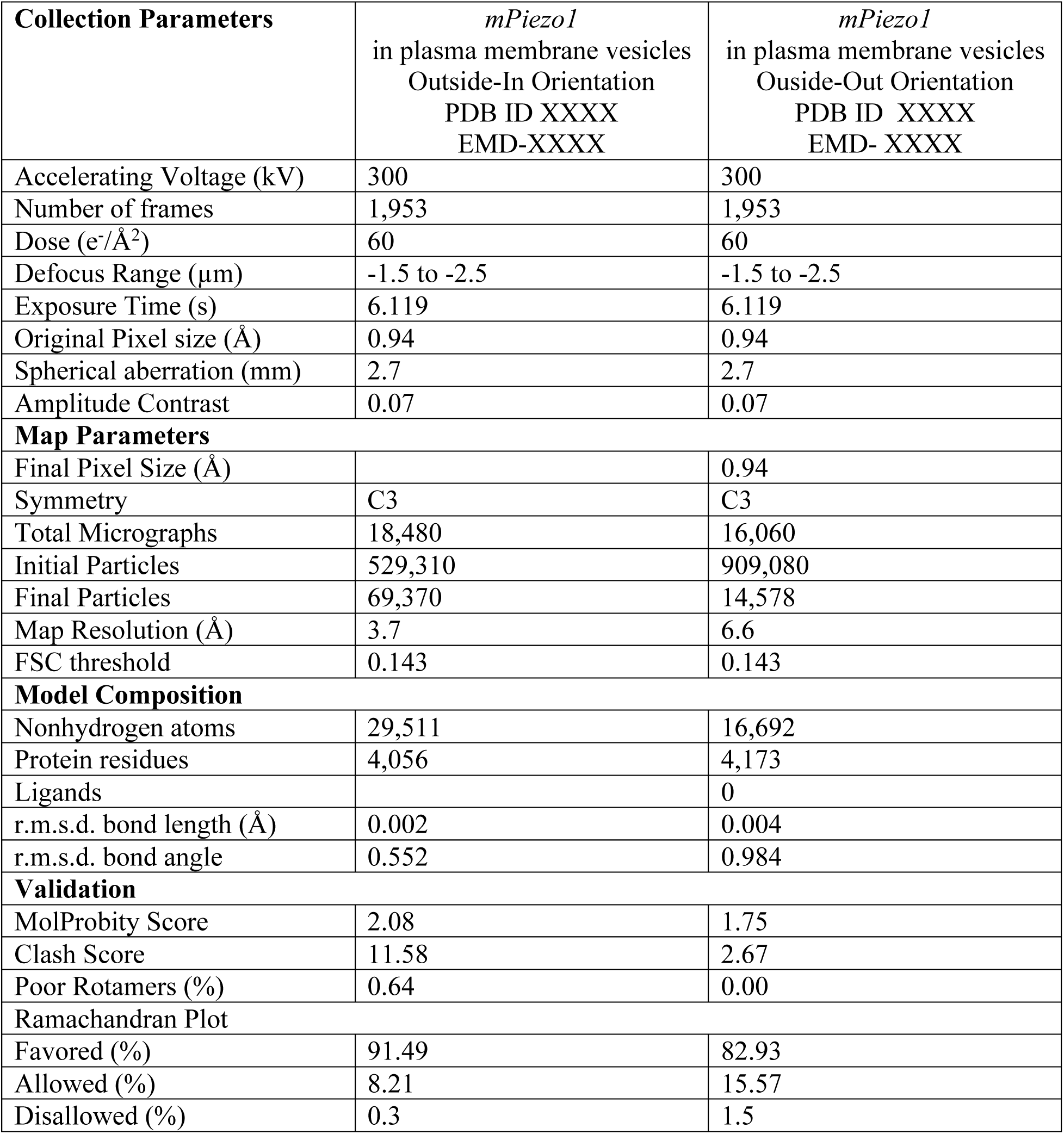
Cryo-EM data collection, map parameters and validation statistics.

